# Re-investigating the coughing rat model of pertussis to understand *Bordetella pertussis* pathogenesis

**DOI:** 10.1101/2021.04.02.438291

**Authors:** Jesse M. Hall, Jason Kang, Sophia M. Kenney, Ting Y. Wong, Graham J. Bitzer, Claire O. Kelly, Caleb A. Kisamore, Dylan T. Boehm, Megan A. DeJong, M. Allison Wolf, Emel Sen-Kilic, Alexander M. Horspool, Justin R Bevere, Mariette Barbier, F. Heath Damron

## Abstract

*Bordetella pertussis* (*Bp*) is a highly contagious bacterium that is the causative agent of whooping cough (pertussis). Currently, acellular pertussis vaccines (aP; DTaP; Tdap) are used to prevent pertussis disease. However, it is clear that the aP vaccine efficacy quickly wanes, resulting in the re-emergence of pertussis. Furthermore, recent work performed by the CDC suggest that current circulating strains are genetically distinct from strains of the past. Emergence of genetically diverging strains combined with waning aP vaccine efficacy call for re-evaluation of current animal models of pertussis. In this study, we used the rat model of pertussis to compare two genetically divergent strains Tohama 1 and D420. We intranasally challenged seven-week-old Sprague-Dawley rats with 10^8^ viable Tohama 1 and D420 and measured the hallmark signs/symptoms of *Bp* infection such as neutrophilia, pulmonary inflammation, and paroxysmal cough using whole body plethysmography. Onset of cough occurred between 2-4 days after *Bp* challenge averaging five coughs per fifteen minutes, with peak coughing occurring at day eight post infection averaging upward of thirteen coughs per fifteen minutes. However, we observed an increase of coughs in rats infected with clinical isolate D420 through 12 days post challenge. The rats exhibited increased bronchial restriction following *Bp* infection. Histology of the lung and flow cytometry confirm both cellular infiltration and pulmonary inflammation. D420 infection induced higher production of anti-*Bp* IgM antibodies compared to Tohama 1 infection. The coughing rat model provides a way of characterizing disease manifestation differences between *Bp* strains.

## Introduction

Whooping cough (pertussis) is a respiratory disease that is caused by the bacterium *Bordetella pertussis (Bp)*. Pertussis is characterized by severe leukocytosis, bronchopneumonia, hypoglycemia, and paroxysmal cough (1, 2). During the catarrhal stage, *Bp* colonizes the upper respiratory epithelium. Once colonization occurs, *Bp* releases toxins such as pertussis toxin (PT), adenylate cyclase toxin (ACT) as well as others. Following the catarrhal stage is the paroxysmal stage that is characterized by severe coughing episodes. Innate and adaptive immune responses result in clearance of the bacterium allowing for the patient or host to convalescence. Despite extensive research, a full understanding of *Bp* pathogenesis remains elusive for reasons such as: vaccine pressure, lack of predictive models, and strain evolution (3). One of the major reasons for this gap of knowledge is the lack of a suitable animal model that shares common disease symptoms as seen in humans. Besides non-human primates, rats are the only other known model of pertussis that share similar coughing manifestation as seen in humans (4). As of now, the origin or the trigger of cough has yet to be determined (5). The rat model of pertussis offers a potential animal model that can be used to evaluate *Bp* pathogenesis.

In 1938, Hornibrook and Ashburn were the first to report that rats infected with *Bp* induced cough-like paroxysms and bronchopneumonia as seen in humans (6). Hornibrook and Ashburn found that young rats were capable of being infected and the bacterium could be cultured from the lungs (6). Infected rats produced coughs that could be heard from a distance of 20 feet (6). Twenty-six of the thirty-one infected rats that did not succumb to the infection had pathology indicative of inflammation in the lungs, most notably early neutrophil infiltration followed by recruitment of mononuclear cells (6). The subsequent studies utilized intrabronchial inoculation of *Bp* encased in agar beads and confirmed leukocytosis and paroxysmal cough by sound activated tape recorders from 5 to 21 days post challenge (7, 8). Further development of the rat model led to the evaluation of PT negative strains and the evaluation of coughs (9). Strain BP357, which is deficient in PT resulted in low cough induction (9). Further evaluation of the rat model demonstrated leukocytosis, weight loss, and paroxysmal cough in Sprague-Dawley rats during the course of *Bp* infection (10). During the development phase of the acellular pertussis vaccine (aP, DTaP), the rat model was used to test vaccine efficacy of various acellular pertussis formulations and was used to validate protection against the onset of leukocytosis and cough (11). Recently, an intranasal rat model has been used to evaluate bacterial factors responsible for cough in *Bordetella bronchiseptica* (*Bb*) (12). BspR, which is an anti-sigma factor of *Bb*, plays some role in cough induction upon infection (12). Collectively, these studies show that the pertussis rat model can be used to critically evaluate *Bp* pathogenesis and disease progression.

In the 1940s, whole-cell pertussis (wP and DTP) vaccines were introduced to protect against *Bp* infection (13). Widespread use of DTP in the United States led to a 90% decrease in the number of reported *Bp* infections (13). Despite the efficacy of this vaccine, serious adverse side-effects ensued, leading to the development of acellular pertussis vaccines. The antigens that are included in the current aPs contain: filamentous hemagglutinin (FHA), fimbriae (FIM), PT, and pertactin (PRN). After the switch from the wP to the aPs, there has been a significant increase in the number of pertussis cases in the US and Europe (14). While nationwide vaccine coverage is 95% in the United States, the incidence of *Bp* infections rose in the past 10 years as the number of aP vaccinated only population has increased in size. (15). While vaccine coverage remains high, the population dynamics are changing, and more people are aP immunized as the new generations are born and as the wP only generations age. Numerous studies have demonstrated waning efficacy of aPs in parallel with the emergence of genetically divergent strains of *Bp* (3, 16–21). Increased surveillance of *Bp* has led to the identification of clinical isolates that do not express PRN, FIM, and even PT the hallmark toxin of the organism (22). One plausible hypothesis to explain these observations is that circulating *Bp* strains have evolved due to acellular vaccine pressure. While we know the current strains are genetically different, we do not know if this genetic variability affects virulence, disease burden, toxicity, or fitness of the pathogen (3).

Early rat studies were performed with intranasal administration of *Bp* for infection (6). Subsequent rat challenge studies used *Bp* encased in agar beads for intrabronchial instillation (7–11, 23). While the agar bead infection method was successful at establishing infection, we aimed to utilize the simplicity of intranasal administration. We sought to re-investigate the coughing rat model of pertussis to compare the pathogenesis of reference strain Tohama 1 to the recent clinical isolate D420, which has been extensively studied in the baboon model of pertussis (24–30). Tohama 1 was first isolated from a case of whooping cough in 1954, while D420 was isolated in 2002 from a critically ill infant in Texas (25, 31). Although Tohama 1 is now a reference strain that has been widely used since Sato and Sato developed the aP but recent data has shown that this strain is an considered an outlier (32, 33). Tohama 1 does expresses PRN but has lower expression of PT and ACT (34). When Tohama 1 was used as a challenge strain in baboons, the baboons did not exhibit symptoms of pertussis despite being infected (26). This led to the selection of recent clinical isolate D420 as the baboon challenge strain, which readily infected and caused disease (26). With 48 years of potential genetic divergence, we sought to understand the differences in pathogenesis between these two commonly studied strains. D420 is known to infect mice and baboons and it belongs to the clade (CDC013) of strains that represented 50% of isolates recovered in the US in 2000 (3). However, it is important to note that D420 has an intact pertactin gene and it does express the PRN protein as confirmed by shotgun proteomic analysis (data not shown).

In our current study, we aimed to re-establish an upper respiratory tract infection model in rats following intranasal challenge with Tohama I and D420. We hypothesized that recent isolate D420 would induce a more severe disease profile compared to Tohama 1, as previous research investigating pathogenesis in rhesus macaques noted that Tohama 1 infected animals did not exhibit overt disease symptoms (26). Sprague-Dawley rats were intranasally challenged with *Bp* and we characterized their disease progression profile over a twelve-day infection. Cough was critically assessed utilizing whole body plethysmography (WBP). Bacterial colonization, leukocytosis, and serological responses were measured as a result of infection. Rats challenged with D420 had increased coughing, greater bacterial burden in the respiratory tract, and a more robust IgM antibody response compared to Tohama 1. The coughing rat model of pertussis can shed light on the pathogenesis of *Bp* and will likely be useful tool for pertussis vaccine evaluation.

## Results

### D420 infected rats have increase number of coughs compared to strain Tohama 1 over the entire course of infection

Over the past decade it is became clear that improvements to the acellular pertussis vaccine strategy are needed due to the rise of PRN mutants, genomic divergence, and epidemiological data. We believe in order to improve pertussis vaccines we need to better understand the pathogenesis of *Bp* in an animal model that shares similar clinical manifestation of pertussis as seen in humans. Here, we intranasally infected seven-week-old Sprague Dawley rats with 10^8^ CFU of *Bp* strain D420 or Tohama 1 (**Fig. 1**) in an effort establish infection and observe *Bp* induced cough. For negative controls (no bacterial challenge), rats were intranasally administered sterile phosphate buffer saline (PBS). Paroxysmal cough is the hallmark symptom of pertussis and is thought to play a major role in transmission of the organism to a new host. To quantify respiratory function during infection, we utilized whole body plethysmography (WBP). WBP instrumentation consists of specialized containment chambers which monitor box flow, temperature, and airflow changes to measure respiratory function. Counting coughs via WBP provides an extremely accurate and unbiased way of counting coughs based on cough waveforms. Rat containment chambers were placed inside a laminar flow hood and connected to a computer that would monitor the rats breathing (**Fig. S1**). Early studies reported that *Bp* infection in young rats induced coughs that were audible by ear (6) and in our preliminary studies this was apparent to us as well (data not shown). Studies in the 1990s, of paroxysmal cough in rats were quantified with an analog sound recording device (7). We hypothesized that rats infected with recent isolate D420 would induce more coughs than reference strain Tohama 1, because rhesus macaques infected with Tohama 1 did not exhibit overt disease symptoms but D420 induced robust coughing in baboons (26). *Bp* infected rats developed cough at days 2-11 (**Fig. 2B-C**), while the mock infected rats (**Fig. 2A**) only had a few isolated coughs, unrelated to infection. Days 1-3 post infection, the average cough count of rats infected with Tohama 1 or D420 was less then 5 coughs per fifteen minutes. The average cough count doubled by day 7 post infection with peak coughing occurring at day 8 post infection with an average of 13 coughs per fifteen minutes of monitoring. After day 8, the average cough count for rats infected with D420 remained above 10 coughs per fifteen minutes, while the rats infected with Tohama 1 averaged less then 5 coughs. To summarize the cough data, the average cough count over the course of infection is shown **(Fig. 2D)**. We observed a significant increase in the number of coughs at day eight post infection of *Bp* infected rats compared to the mock challenge control. Rats infected with D420 coughed a total of 949 times, whereas rats infected with Tohama 1 coughed a total of 724 times over the entire twelve-day infection. To analyze differences in cough count between strains, we utilized Area Under the Curve (AUC) analysis **(Fig. 2E)**. AUC analysis allows us to quantify the number of coughs over the entire course of infection per rat. Upon this analysis, we note a significant increase in the number of coughs for rats infected with D420 compared to the mock challenged rats over the twelve-day infection, while there was no significant difference observed in Tohama 1 infected rats compared to mock challenge (non-infected).

**FIG 1.**
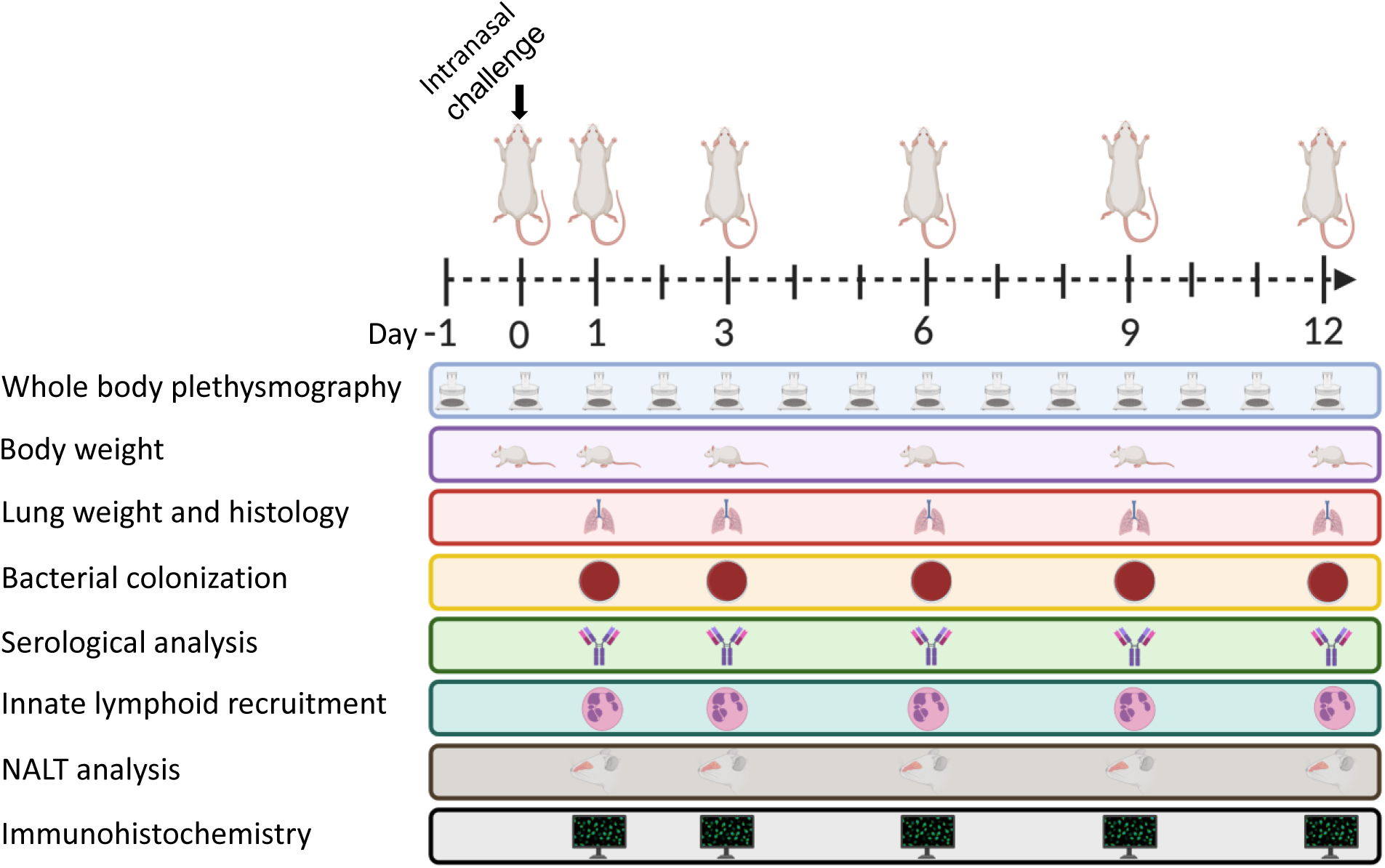
Experimental design of intranasal infection of *Bp*. Schematic representation of Sprague-Dawley rats intranasally infected with *Bp*. Whole body plethysmography was used to measure cough and analyze lung function over the course of a twelve-day infection. Body weight, Bacterial burden of the respiratory tract, innate lymphoid recruitment, and antibody titers were measured at the days 1, 3, 6, 9, and 12 post challenge. The left lobe of the lung and NALT were sectioned and stained with hematoxylin and eosin for histological analysis and immunohistochemistry was performed to visualize bacteria in the left lobe of the lung and nasal cavity.

**FIG 2.**
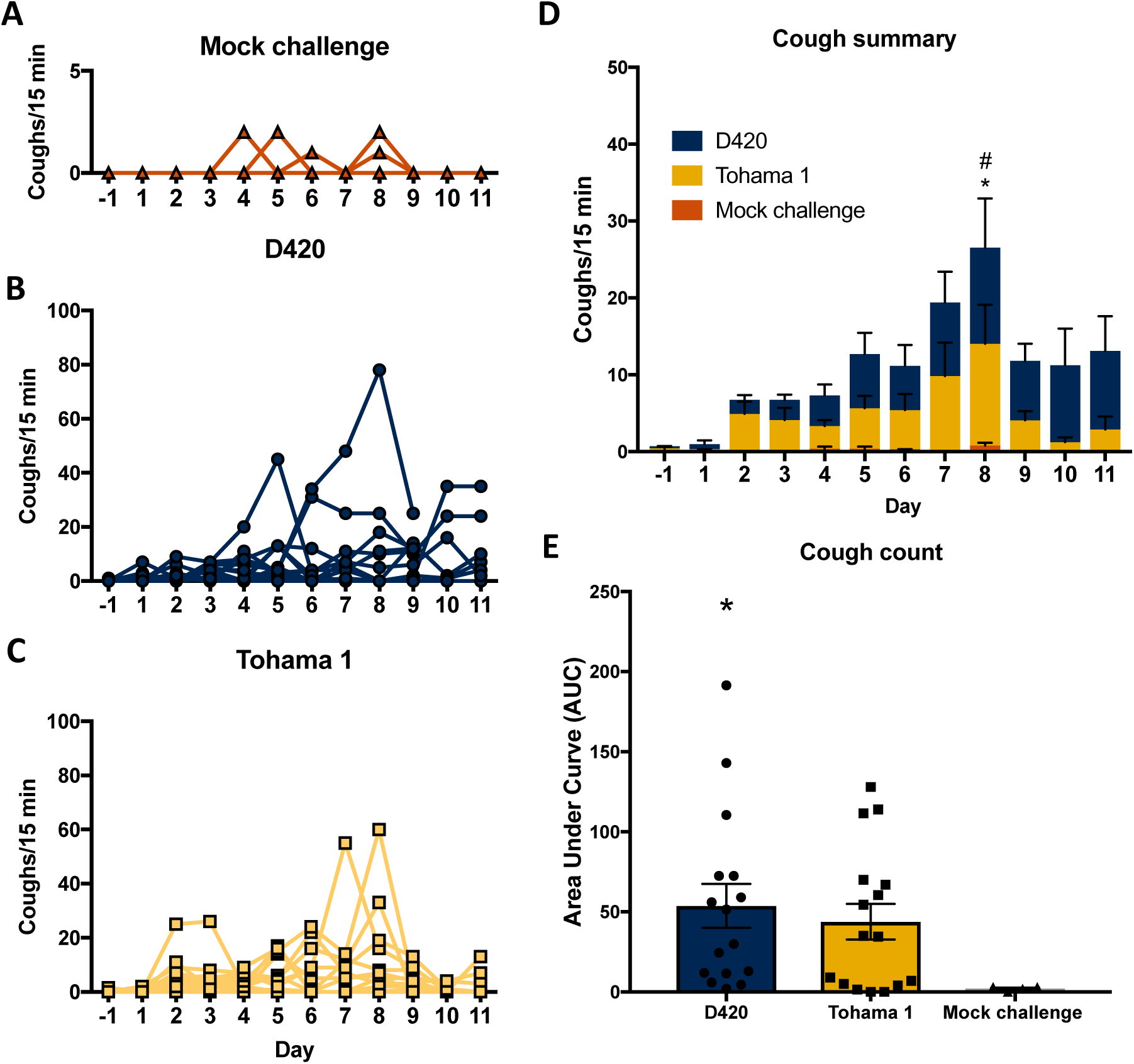
Infection of *Bordetella pertussis* induces cough in Sprague-Dawley rats. Coughs were measured each day of the twelve-day infection using whole body plethysmography. Coughs were counted for **(A)** mock challenge rats, **(B)** rats infected with D420, and **(C)** rats infected with Tohama 1. **(D**) Summary of results (**A-C)** shown as mean ± SEM (*n* = 4-8). **(E)** Area under the curve is represented as averages of coughs per fifteen minutes for each rat. *P* values were determined by two-way ANOVA followed by a Bonferroni comparison test and unpaired Student t-test for AUC, \**P <* 0.05 D420 compared to the mock challenged control group, *^#^P* < 0.05 Tohama 1 compared to the mock challenged control group.

### *Bp* infection causes pulmonary distress

Infection with *Bp* leads to mucus production, lung damage, and invasion of cellular infiltrates into the bronchioles of the lung (35). The lungs of infants with *Bp* exhibit edema, necrotizing bronchiolitis, and inflammation in the lung leading to respiratory distress (35). WBP was used to quantify the pulmonary distress over the course of infection. Pulmonary distress was measured by calculating the enhanced pause (PenH) of the animal (**Fig. 3A-B**). The higher the PenH value, the increased respiratory distress of the animal. Compared to the mock challenged rats both D420 and Tohama 1 infected rats had an increase in PenH (**Fig. 3A-B**). We observed a significant increase at day 9 with rats infected with D420 and at day 6 with rats infected with Tohama 1 compared to mock challenge.

**FIG 3.**
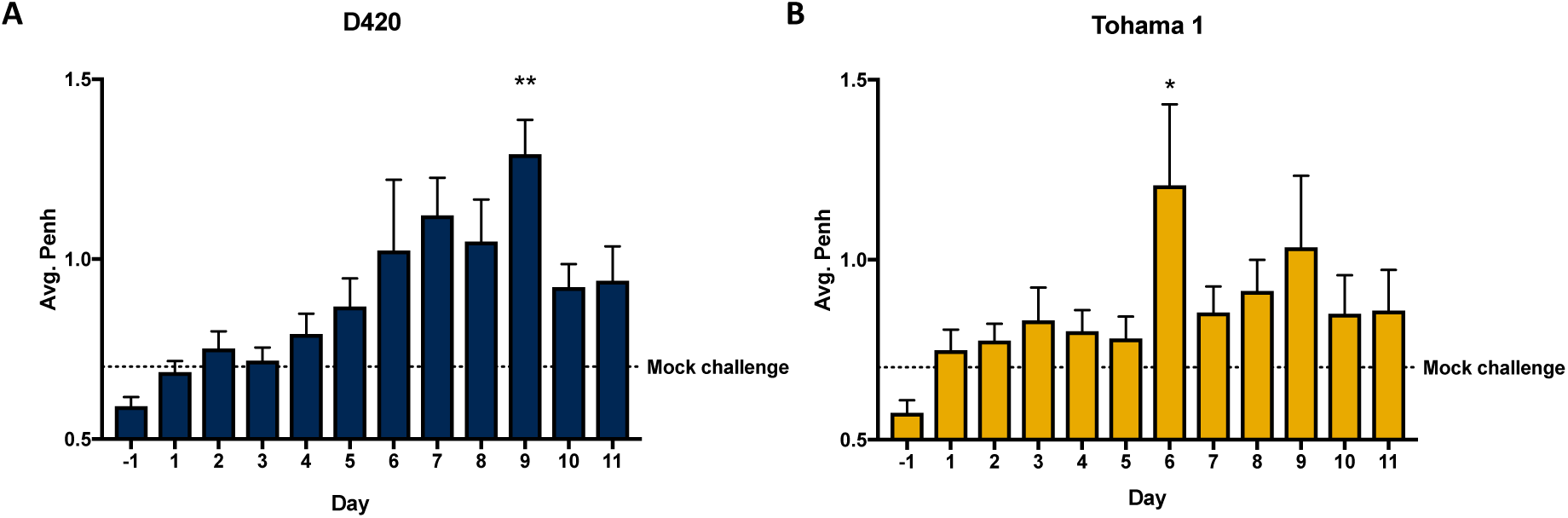
*Bordetella pertussis* infection impairs respiratory capacity. Over the course of infection, Sprague-Dawley rat’s respiratory function was analyzed using whole body plethysmography. Each day at 5p.m., rat’s bronchial restriction was determined by utilizing WBP by calculating for PenH. Penh was calculated **for (A)** D420 and **(B)** Tohama 1. Results shown as mean ± SEM (*n* = 4- 8). *P* values were determined by two-way ANOVA, \**P <* 0.05, \*\**P <* 0.01 compared to the mock challenge group followed by Dunnett’s comparison test.

### Histological assessment of the lung and nasal cavity

To further evaluate the inflammation in the lung, we utilized histology to confirm both acute and chronic inflammation in response to *Bp* infection. At days 1, 3, 6, 9, and 12 post challenge the left lobes were collected, sectioned, and stained with hematoxylin and eosin (H&E) (**Fig. 4A&C**). Whole-lung images were taken to assess cellular infiltration (**Fig. S2A&B**). Once stained, the blinded slides were then scored by a board-certified pathologist. Mock challenge animals exhibited minimal inflammatory infiltrates consisting of focal accumulations of mononuclear cells in the parenchyma and occasional infiltrates of neutrophils surrounding blood vessels (**Fig. 4A**). Rats infected with Tohama 1 and D420 had significant increase in their acute inflammation scores compared to mock challenge animals at day 1 and 3 post infection (**Fig. 4B**). Day 1 post infection had the highest acute inflammation score (**Fig. 4B**). Rats exhibited mild to moderate neutrophil infiltration of the parenchyma, blood vessels, and the bronchioles (**Fig. 4A**). Markers of acute inflammation resolved after day 6 post challenge; however, infected rats at day 9 exhibited obvious differences in the lung associated with mild to moderate infiltration of mononuclear cells and higher chronic inflammation compared to mock challenge animals (**Fig. 4C**). Interestingly, rats infected with D420 had significant higher chronic inflammation score compared to rats infected with Tohama 1 (**Fig. 4D**). At day 12 post infection, we notice moderate resolution of inflammation in both D420 and Tohama 1 infected rats (**Fig. 4B&D**). We also observed a significant increase in lung weight, which can also be associated with elevated inflammation, at days 1 and 9 post challenge in D420 infected rats compared to the mock challenge control (**Fig. S3A**). At day 12 post challenge rats infected with Tohama 1 had a significant increase in lung weight compared to the mock challenge rats (**Fig. S3A**). These results confirm that *Bp* infection induces inflammation in the lung of coughing Sprague-Dawley rats.

**FIG 4.**
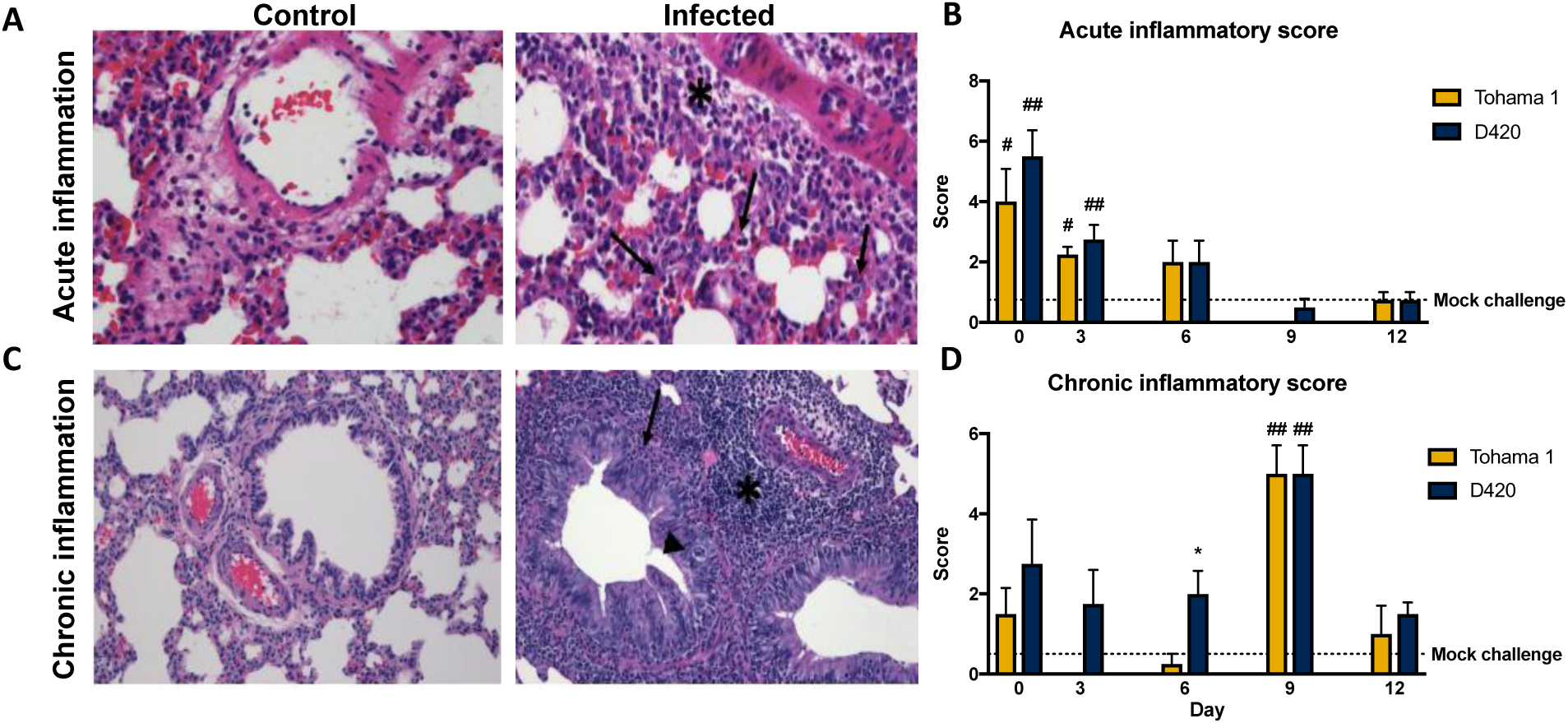
*Bordetella pertussis* infection induces acute and chronic inflammation in the lung of Sprague-Dawley rats. The left lobe of the lung was sectioned and stained with hematoxylin and eosin from rats infected with D420, Tohama 1, or the PBS control. **(A)** Representative images of acute inflammation demonstrating neutrophil recruitment around surrounding blood vessel (asterisk) and parenchyma (arrows) at 400x magnification. **(B)** Average acute inflammatory score of the lung by the predominance of neutrophils in the parenchyma, blood vessels, and airways. **(C)** Representative images of chronic inflammation showing mononuclear cells surrounding the blood vessel (asterisk), lamina propria (arrow), and bronchioles (arrowhead) at 200x magnification. **(D)** Average chronic inflammatory score of the lung characterized by mononuclear infiltrates in the parenchyma, blood vessels, and airway. Histological assessment was determined blinded with no knowledge of the treatment groups. Results are shown as mean ± SEM (*n* = 4) *P* values were determined by one-way ANOVA followed by Dunnett’s comparison test, \**P <* 0.05 compared between challenge groups. ^#^*P <* 0.05, ^##^*P <* 0.01 compared to mock challenge.

Due to the fact that *Bp* is a mucosal pathogen, we sought out to investigate any phenotypic changes or inflammation in the Nasal-Associated Lymphoid Tissue (NALT) of pertussis infected rats via histology. NALT helps elicit immunity against airborne and mucosal pathogens (36). The NALT is located above the hard palate and studies have also shown that with the introduction of antigen via vaccination or bacterial components, cellular expansion of the NALT ensues (37–41). The NALT was stained with H&E, and we observed an increase in cellular infiltrate and a size enlargement in the NALTs of *Bp* infected rats compared to the mock challenged (**Fig. S4A**). ImageJ analysis was used to measure the area of both the left and right NALT of each animal (**Fig. S4B**). We observed a significant increase in the area of the NALT of rats infected with D420 at day 6 and 9 compared to mock challenge, while we only saw an increase in the NALT area in Tohama 1 infected rats at day 9 post challenge compared to our mock challenge control (**Fig. S4B**). This data could mark the potential cellular expansion of the NALT upon infection with *Bp*. Further analysis of the expanding cell populations in the NALT will warrant more analysis especially in the context of vaccine immunity.

### D420 infected rats have increased bacterial burden in the respiratory tract

We determined bacterial burden in the respiratory tract for Tohama I and D420 challenged rats at days 1, 3, 6, 9, and 12. Over the coughing timeline, we see an overall decrease in viable bacteria in the lung, trachea, and nasal cavity. In the lung, we observed a significant increase in the recovered bacteria in the D420 infected rats compared to the Tohama 1 infected rats at both days 3 and 6 (**Fig. 5A**). The same trend is also seen in the nasal cavity at day 1 post infection (**Fig. 5C**). We observed no difference in the viable bacteria recovered in the trachea (**Fig. 5B**). Previous pertussis studies utilizing the rat model of pertussis noted weight loss from intrabronchial infection of *Bp* strain 18-323 (10). *Bp* infection is sufficient to cause weight loss in rats (10). To confirm this using intranasal infection, body weight was measured before challenge and upon euthanasia to calculate percent weight change. We observed a significant decrease in weight gain in rats infected with *Bp* strain D420 and Tohama 1 compared to the mock challenged at day 12 post infection (**Fig. S3B-D**). Next, we aimed to compare the bacterial colonization between outbred mice and rats in an effort to relatively compare the bacterial burden between models. We utilized the bacterial burden data in CD1 mice and compared them to our data in rats (42). To address the body weight differences between the mice and rats, we used the CFUs/organ and divided them by their respective body weight for a crude comparison. Our data suggests that despite having a higher initial challenge dose in the rat, the bacterial burden remaining in the respiratory tract is higher in a mouse compared to the rat later during infection (**Fig. S5**). In general, the rat model is likely a lower challenge dose per body weight model of pertussis compared to mouse models.

**FIG 5.**
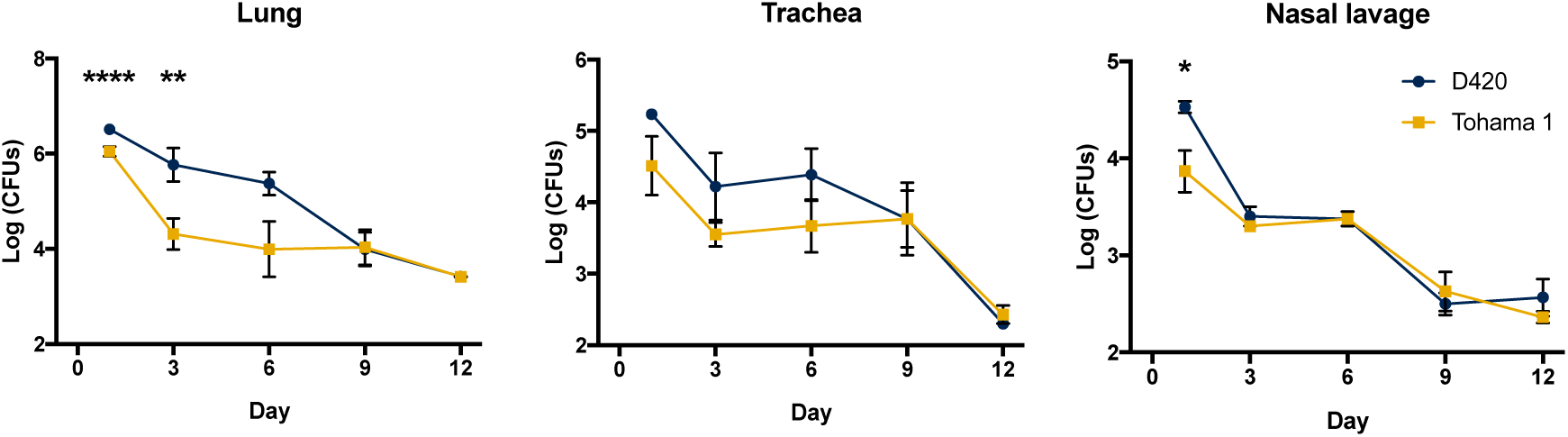
Analysis of bacterial burden over the course of *Bp* infection. Bacteria were quantified by serially diluted CFUs following intranasal challenge. CFU counts were determined from **(A)** lung homogenate **(B)** trachea and **(C)** nasal lavage. Results are shown as mean ± SEM (*n* = 4) *P* values were determined by two-way ANOVA corrected with Bonferroni comparison test, \**P <* 0.05, \*\**P <* 0.01, \*\*\*\**P <* 0.0001 compared between D420 and Tohama 1.

### Immunofluorescence confirms colonization of *Bp* in the lungs and nasal cavity

We next wanted to assess the location of bacterial colonization in the nasal cavity and visualize any bacteria left behind following flushing of the nares. We sectioned the skulls after flushing the nasal cavity and utilized Immunofluorescence (IF). After visually scanning the nasal cavity with confocal microscopy, we found bacteria in the nasal turbinates and the NALT (**Fig. 6A&C**). We also confirmed that the bacteria were trapped in the crypts between the epithelial cells (**Fig. 6B**). The relative locations of where the bacteria were found in the nasal cavity can be seen in (**Fig. S6A**). Our results show that *Bp* remained in the nasal cavity over the entire course of infection even after flushing the nares and the bacteria also remained captured in the cilia of the nasal turbinates (**Fig. S6B**). The flushing of the nasal cavity underrepresents the total number of *Bp* in the nasal cavity. In the lung, *Bp* colonizes the bronchioles, while the alveolar spaces remain non-colonized (**Fig. 6D&F**). IF was performed on the nasal cavity and lung of mock challenge animals to access non-specific binding (**Fig. S7**). To assess any potential differences between the strains we blindly manually counted the labeled microcolonies in the lung and nasal cavity. It is also worthy to note, we found no differences in microcolony counts in the lung or nasal cavity between the strains (**Fig. S6 C-D**) despite our observed differences in bacterial burden by CFU enumeration. CFU is likely a better indicator of overall bacterial burden, but IF revealed that *Bp* is hiding and well attached in distinct locations in the airway.

**FIG 6.**
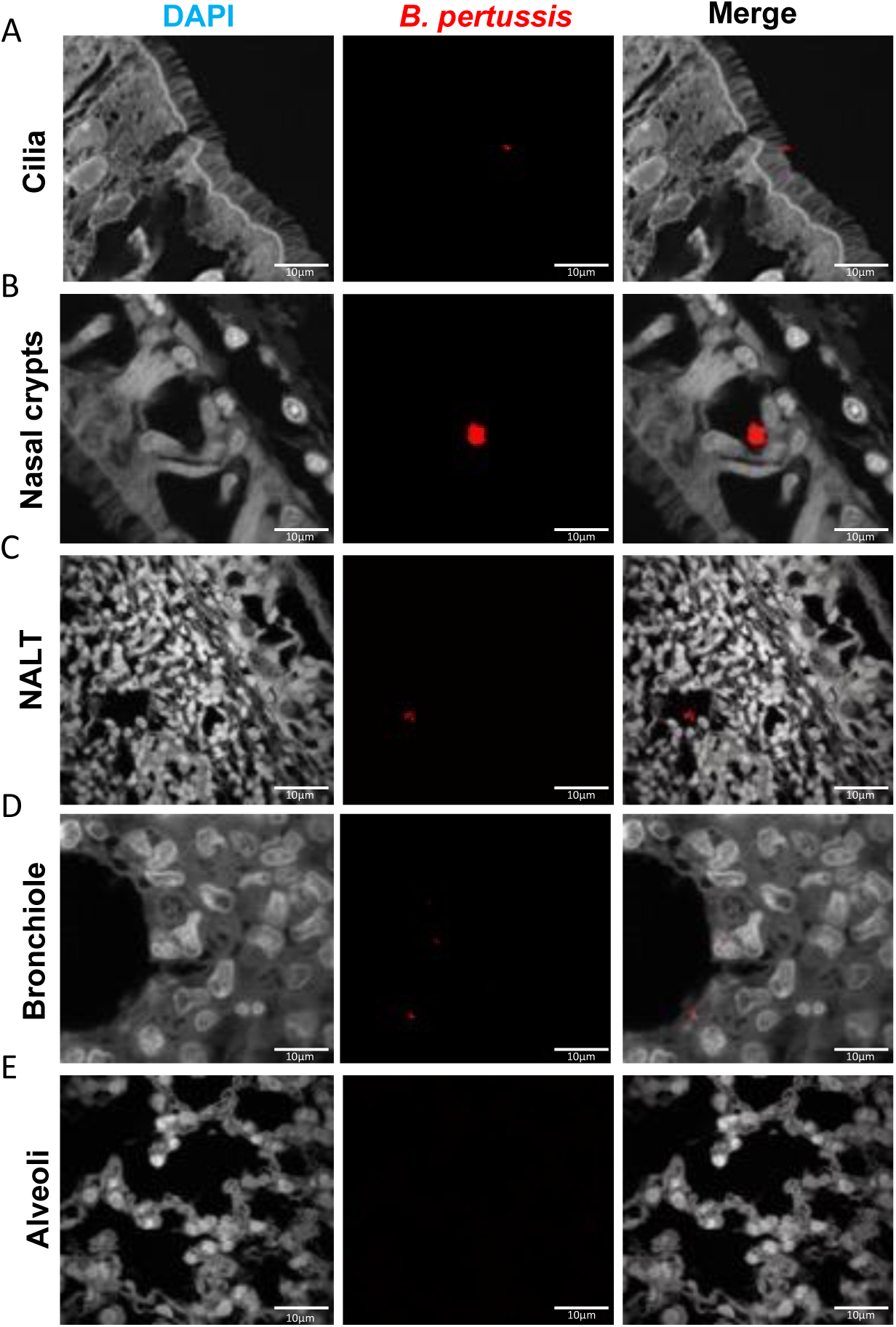
Immunofluorescence (IF) staining of *Bp* localization in the respiratory tract. *Bp* was labeled using a polyclonal antibody to FHA and counter tagged with a fluorescently conjugated antibody (Texas-Red). Sections were counterstained with DAPI. **(A-C)** Representative images of *Bp* in the nasal cavity over the course of infection. *Bp* was found captured in the cilia of the nasal cavity as well as the NALT. **(D-E)** Representative images of the bronchiole and alveoli of infected rats. *Bp* was found localized in the bronchioles over the course of infection and absent in the alveoli.

### Measurement of recruitment of neutrophils in response to *Bp* infection

Previous studies utilizing the rat model of pertussis noted leukocytosis and an increase in the amount of total white blood cells using agar-encased *Bp* for infection (8). Neutrophils circulating through the blood were measured during intranasal infection by antibody staining and flow cytometry, as *Bp* infection results in varying amounts of neutrophilia (35, 43). The percentage of CD45+ neutrophils in the blood were increased of infected rats compared to the mock challenged animals (**Fig. 7A**). Specifically, a significant increase in the number of circulating neutrophils in the blood of rats infected with D420 at days 1 and 9, with Tohama 1 infected rats only having a significant increase to mock challenge at day 9 (**Fig. 7A**). Next, cytokines concentrations in the lung and serum were measured to identify factors that could contribute to the recruitment of neutrophils, specifically IL-6 and IL-17. IL-6 is a potent inducer of Th17 polarization and been found elevated during murine infection with *Bp* (42, 44–46). Surprisingly, we did not observe any significant changes in IL-6 due to *Bp* infection (**Fig. 7B&C**). It has been well documented in the mouse model that the production of IL-17 in pertussis plays a role in the increase of circulating neutrophils (47). There were no significant changes in IL-17a in the serum or lungs of *Bp* infected rats compared to mock challenge (**Fig. 7D&E**). Th1 and Th2 cytokines and chemokines in the lung and serum were also measured (**Fig. S7-Fig. S10**). We did not observe an increase in the induction of cytokines by infection overall; however in the serum, rats infected with D420 had a significant increase in IL-5 compared to Tohama 1 infected rats at day 12 post infection (**Fig. S8)**. There was a significant increase in IL-5 and TNF-alpha at day 6 post infection in the lung in rats infected with Tohama 1 compared to D420 (**Fig. S9**). We also observed significant increases in chemokines MIP-1a, RANTES, MCP-3, MIP-2, and IP10 in rats infected with D420 at day 12 post challenge in the serum compared to rats infected with Tohama 1 (**Fig. S10**). In the lung, MIP-2 and IP10 were increased in rats infected with D420 compared to mock challenge control (**Fig. S11**). Despite slight differences highlighted above we did not observe a massive difference in the cytokine and chemokine responses during infection.

**FIG 7.**
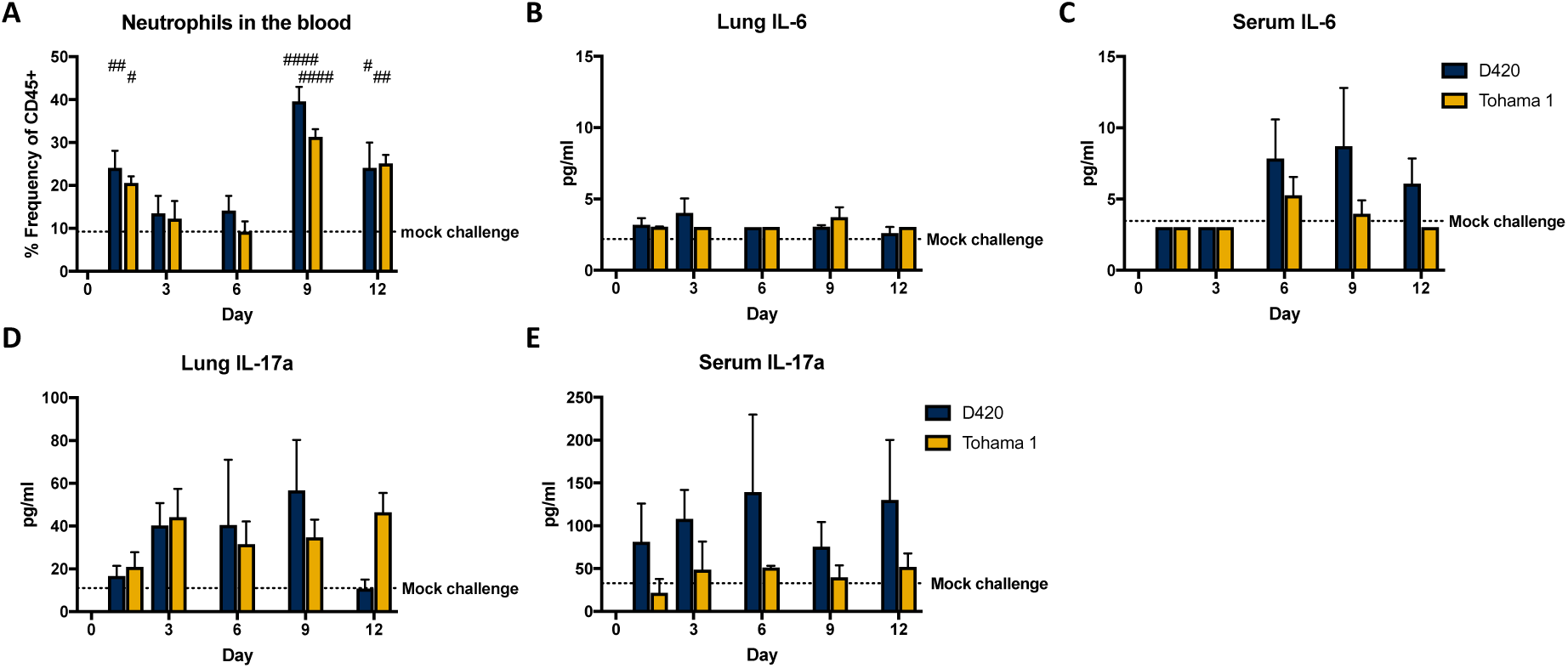
Neutrophil recruitment and proinflammatory cytokine production after challenge determined by flow cytometry and multiplex immunoassay analysis. **(A)** Blood neutrophil count of Sprague-Dawley rats infected with *Bp*. At days 1, 3, 6, 9 and 12 post infection, rats were euthanized, and the blood was collected via cardiac puncture. Flow cytometry was used to quantify the amount of neutrophils (CD45^+^ CD161^-^ B220^-^ CD43^+^ His48^hi^) in the blood. Quantification of the percentage of single, CD45+ cells. Dotted line represents the average frequency of CD45+ neutrophils in the mock challenge animals over the course of the study. **(B-C)** Analysis of cytokine IL-6 from supernatant of lung homogenate and serum. **(D-E)** analysis of cytokine IL-17 in the serum and lung homogenate supernatant. Cytokines analyzed using ProcartaPlex multiplex immunoassay kit. Results shown as mean ± SEM (*n* = 4). Dotted line represents the average mock challenged cytokine levels measured. *P* values were determined by one-way ANOVA followed by Tukey comparison test, ^#^*P <* 0.05, ^##^*P <* 0.01, ^####^*P <* 0.0001 compared to mock challenge.

### Serological analysis of *Bp* specific antibodies

Enzyme-linked immunosorbent assays (ELISA) were performed to determine the serological response generated from *Bp* infection. We measured IgM and IgG antibody titers in the serum to whole bacterium and *Bp* associated virulence factors at days 1, 3, 6, 9, and 12 post infection. In previous studies utilizing the rat model, intrabronchial infection induced the generation of IgG antibody titers to sonicated *Bp* 28 days post infection (8). Intranasal infection of strain D420 has generated significant anti-PT IgG antibody titers in the baboon model at days 17-19 of infection (26). However, analysis of antibody titers generated against an intranasal pertussis infection in the rat has yet to be determined and we wanted to characterize the primary antibody response to *Bp*. We observed a significant 10-fold increase in the anti-*Bp* IgM antibody response at day 9 and an almost 20-fold increase 12 days post challenge in rats infected with D420 compared to mock challenge, while we also saw a significant 6-fold increase in anti-*Bp* IgM at day 12 post infection to the amount of anti-*Bp* IgM from Tohama 1 infection (**Fig. 8A&B**). At day 12 post infection, we observed that *Bp* infection induced significant production of anti-*Bp* IgG in the serum compared to mock challenge animals (**Fig. 8C&D**). At day 12 post infection, Tohama 1 infected rats had a significant increase, nearly double anti-*Bp* IgG antibody titers compared to rats infected with D420. AUC analysis revealed a significant increase in anti-*Bp* IgM antibodies from rats infected with D420 compared to Tohama 1 infected rats (**Fig. S12A**). However, no difference was observed in AUC of the anti-*Bp* IgG antibody response over the total course of infection between the bacterial challenge groups (**Fig. S12B**). It is also important to note we also did not see a significant increase in the generation of antibodies titers to PT, ACT, PRN, and FHA in *Bp* infected rats compared to mock challenge infected animals (**Fig. 8A-D**). Graphs representing statistical significance of the antibody titers between D420 and Tohama 1 infected rats can be observed in the supplementary data (**Fig. S13**).

**FIG 8.**
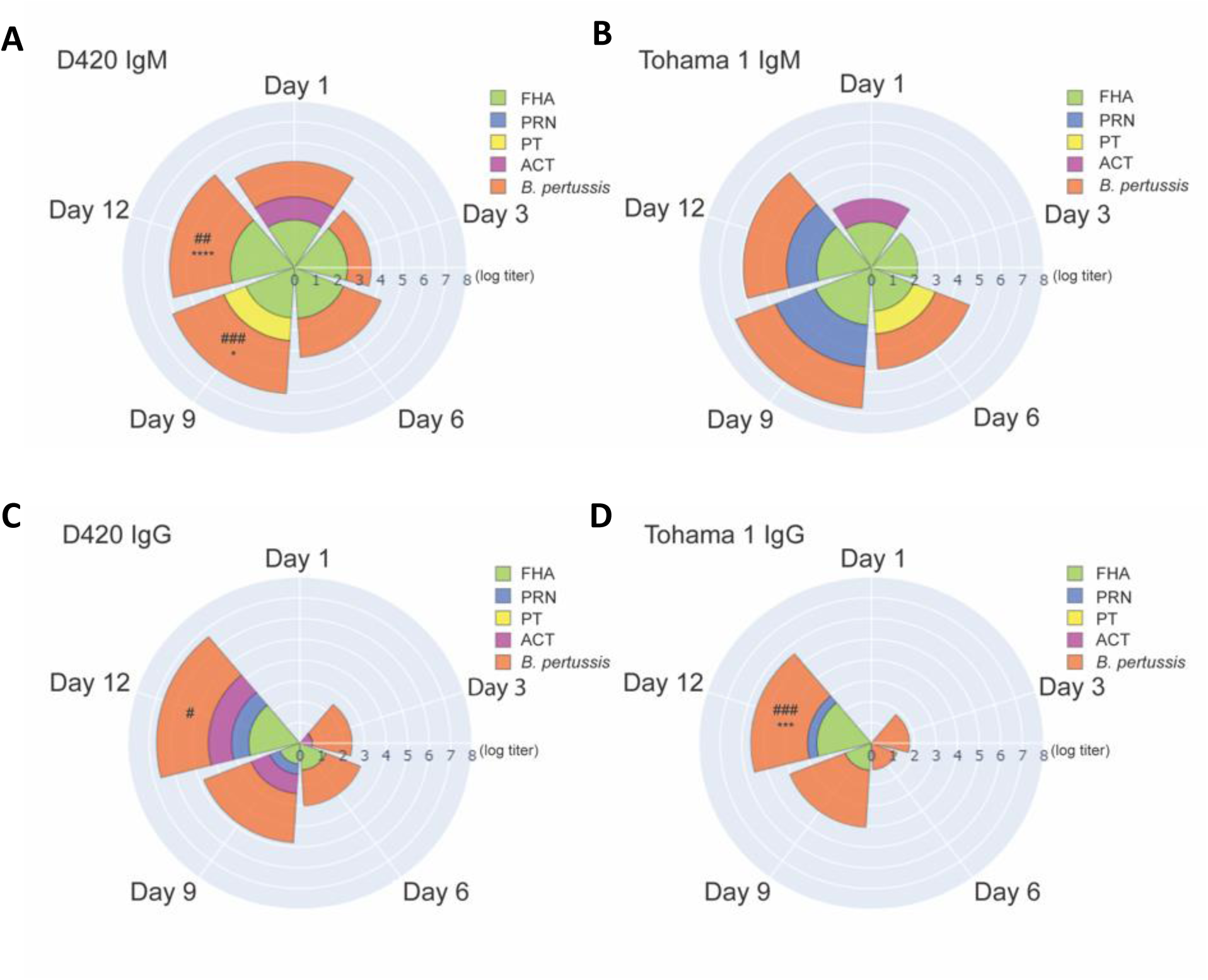
Measurement of serum antibody titers over the course of *Bp* infection. ELISA was used to compare serological responses from rats IN challenge with *Bp*. Total IgM (A-B) and IgG (C-D) serum antibody titers were measured from challenged rats against *Bp*, FHA, PRN, PT, and ACT. (*n* = 4). *P* values were determined by two-way ANOVA corrected with Bonferroni comparison test, \**P <* 0.05, \*\*\**P <* 0.001, \*\*\*\**P <* 0.0001 compared between infection groups. For comparison to mock challenged control group one-way ANOVA was used followed by Tukey comparison test, *^#^P <* 0.05, *^##^P* < 0.01, ^###^*P <* 0.001.

## Discussion

To date, the only known reservoir for *Bp* is humans, making the development of a suitable animal model challenging. Multiple models have been studied to recapitulate similar disease symptoms seen in humans, as well as provide insight into the pathogenesis of *Bp*. We set out to re-investigate the rat model of pertussis that was originally introduced in 1938 (6). In our study, we have confirmed that intranasal administration of *Bp* led to bacterial colonization of the upper and lower respiratory tract (**Fig. 5**). Through intranasal instillation, bacteria were able to infect the nasal cavity, trachea, and lung post challenge. This is important to note as infants with pertussis experience pneumonia more frequently than adolescents and adults (35). Mouse models of pertussis also rely on depositing large numbers of bacteria into the lung to establish infection (2). Here, our data shows that through intranasal infection, we establish an infection of both the upper and lower respiratory tract, resembling the infection seen in adolescents and adults with pertussis. Previous reports have shown *Bp* infection of rats induced paroxysmal coughs, and these coughs were counted by analog audio recordings (7). Utilizing WBP we are able to precisely and digitally count coughs and measure respiratory capacity for the first time with rats infected with *Bp* (**Fig. 2&3**). Here, coughs induced from infection appeared at day 1-2 post challenge, with the number of coughs gradually increasing with peak cough occurring at day 8 post challenge. Rats challenged with intrabronchial instillation of *Bp* noted similar peak cough counts approximately 10 days post challenge (48). This is also one of the first studies utilizing WBP to count coughs in rats with bacterial infection. We did observe some infected rats that did not overtly cough. This reflects the frequency of coughs in adolescents and adults (49). Previous studies using baboons also show that *Bp* infection elicits severe coughs lasting over 2 weeks post challenge with peak coughs per hour being at day 4 post challenge (26). It is also important to note that at peak coughing, the bacterial burden was low, which was also observed in previous *Bp* infected rat research (7). The data suggest that other factors could be playing a role in cough production despite the bacterial infection being resolved. We have also observed that the increased bronchi restriction coincides with the increased coughs during days 6-11 post challenge (**Fig. 3)**. The decreased lung capacity and increase in cough could be further associated with an increase of chronic inflammation in the lung (**Fig. 4**). Our data suggests that the coughing rat model of pertussis can be used to potentially evaluate the mechanism of cough, which still requires further investigation.

Initial studies characterizing the serum antibody response to rats infected with *Bp* noted an increase antibody response to endotoxin correlated with the low amount of cultivable organisms in the lung following intrabronchial challenge (7). Hall et al. demonstrated that rats infected with *Bp* become sera positive to whole cell *Bp* lysate, FHA, and PT at 28 days post intrabronchial challenge (8). Anti-FHA antibody titers were not detected until day 16 post challenge, while anti-PT antibody titers were not detected until day 28 post challenge (23). In our study, we characterized the antibody response following intranasal challenge of *Bp* over the whole infection timeline. Using *Bp* coated ELISA plates, we detected anti-*Bp* IgG antibody titers in rats infected with *Bp* at day 9 post challenge with a significant increase compared to mock challenge observed at day 12 (**Fig. 8**). We also measured a significant increase in anti-*Bp* IgM antibody titers in rats infected with D420 compared to the mock challenge and Tohama 1 infected animals (**Fig. 8**). One potential hypothesis that could explain this observation is the genetic differences between the two strains as D420 harbors the *ptxP3* allele, leading to an increase in PT production (25). PT has been shown to suppress *Bp* antibody responses in the serum and limit expression of antigen presenting receptors (50–53).

With the increased interest in developing new pertussis vaccines, it is imperative for the development of animal models to evaluate the pathogenic potential of current emerging strains, as well as identify any potential differences in disease burden between genetically divergent strains. Sato and Sato first used *Bp* strain Tohama 1 in the development of the acellular pertussis vaccine, and Tohama 1 is still being used in pertussis research today, notably due to its genetic malleability (54). D420 has been instrumental in the development of the baboon model and has also been used to study vaccine efficacy in regards to transmission in the baboon model (29). In our study, we observed multiple differences between Tohama 1 and D420 ranging from coughing, bacterial colonization, and serological responses (**Fig. 2, 5, 8**). We hypothesize that these observed differences could be due to the increased amount of PT through the *ptxP3* allele of D420 (25). Strains, such as D420, harboring the *ptxP3* allele lead to increase in virulence, PT production, and prevalence (55, 56). A previous study utilizing the intranasal mouse model has shown an increase in bacterial colonization with *ptxP3* strains (57). Parton et al. illustrated that infection with a PT deficient strain was unable to induce cough (9). It is possible that there are other factors beyond PT that can account for the observed virulence differences between Tohama I and D420. In the rat model, unlike the mouse, we also did not measure any significant increases in IL-17 and IL-6 in the lung and serum following challenge. IL-17 and IL-6 are increased following *Bp* infection in mice and baboon (30, 44, 58). Further investigation is needed to explain the observed differences.

In summary, we have demonstrated that the rat model of pertussis can be used as a tool to further study *Bp* pathogenesis and recapitulate some of the similar symptoms of pertussis as seen in humans. Pertussis is often described as a 100 day cough, because adolescents and adults extended duration of coughing (59). Coughing episodes in infants with pertussis can lead to vomiting, choking, gagging, and apneic episodes that can lead to seizures (60). Mice have not been used to evaluate coughing manifestation in pertussis until recently, and baboons are expensive and limited to specialized facilities (61). The coughing rat of pertussis provides feasibility through low husbandry costs, availability of animals, and ease of use as a model. In this study, we carefully compared two established strains that have been used in the pertussis field, Tohama 1 and D420. In future studies, we aim to evaluate whole cell and acellular vaccine mediated immunity against D420. Armed with this model, we can further our understanding of pathogenesis, host response during pertussis, genetic divergence between strains, and vaccine-mediated immunity.

## MATERIALS AND METHODS

### *Bordetella pertussis* strains and growth conditions

*Bp* strain Tohama 1 was graciously provided by Dr. Peter Sebo (Czech Academy of Sciences) and strain D420 was acquired from the CDC Pertussis Lab provided by Drs. Maria L. Tondella and Michael Weigand. *Bp* strain Tohama 1 and D420 were cultured on Bordet Gengou (BG) agar (Remel^TM^ Cat. R45232) that was supplemented with 15% defibrinated sheep blood (Hemostat Laboratories Cat. DSB500) (62). Bacteria were cultured on BG plates for 48 hrs at 36^°^C. *Bp* was transferred from BG plates with polyester swabs (Puritan Cat. 22-029-574) and transferred into 20 ml Stainer-Scholte liquid media (SSM) in new 125 ml flasks (Thermo Fisher Scientific Cat. FB500125) (63). The liquid cultures were grown for 24 hrs at 36^°^C inside a shaking incubator at 180 rpm.

### Intranasal challenge with *B. pertussis*

Seven-week-old ∼170g female Sprague-Dawley rats (Charles River Cat. 001CD) were used for challenge. *Bp* was grown as described above. The rats were then anesthetized with ketamine and xylazine 50-100/5-10 mg/kg and challenged with 10^8^ CFUs (100μl intranasally), administering two 50μl doses one in each nostril (**Fig. 1**). Body weight of each rat was recorded before challenge and immediately post euthanasia. Rats were euthanized at day 1, 3, 6, 9, and 12 post bacterial challenge. Mock challenge animals (no bacteria) were administered 100μl of sterile endotoxin free phosphate buffer saline (PBS) (Thermo Fisher Scientific Cat. TMS012A) intranasally. Upon euthanasia blood was collected via cardiac puncture and transferred into ethylenediaminetetraacetic acid (EDTA) (BD Cat. 365974) and serum separation (BD Cat. 026897) tubes. Blood collected in the EDTA tube was used for flow cytometric analysis, while blood collected int the serum separation tubes were used to isolate the serum via centrifugation (15,000 x *g* for 3 min) and used for serological and cytokine analysis. Lung and trachea were excised and homogenized to determine bacterial burden. Upon removal, the wet weight of the lungs was recorded. Lungs were then collected in gentleMACS C tubes (Miltenyi Biotec) and homogenized using gentleMACS™ Dissociator (Cat. 130-095-927). The trachea was homogenized using a Polytron homogenizer. To determine the bacterial burden in the nares, 2mls of sterile 1x PBS was flushed through the nares and collected for plating. Serial dilutions of the homogenates and nasal collection were plated on BG plates supplemented with ceftibuten (Sigma-Aldrich Cat. SML0037) 10 µg/ml to decrease growth of normal rat respiratory tract flora.

### Analysis of coughing and bronchiole restriction using whole-body plethysmography

To quantify respiratory function during infection, we utilized a Buxco® FinePointe^TM^ Whole Body Plethysmograph instrument (WBP) (DSI). FinePointe^TM^ software was used for collecting, analyzing, and reporting the breathing data. Everyday post challenge for twelve consecutive days at 5:00PM the rats were placed inside designated chambers to acclimate for 5 min. We chose this time, not only due to rats being nocturnal, but this was the time the animals were most awake and active as confirmed by video camera (data not shown). Following acclimation, the rat’s respiratory profile was recorded for 15 min. Each chamber is fitted with a transducer that measures the changes in box flow and airflow of the subject. The chambers are also fitted with a screen pneumotach that allows airflow in and out of the chamber that can be recorded. Coughs were counted during the designated 15 min, and enhanced pause (PenH) was calculated to signify bronchiole restriction. Coughs were counted based on large box flow changes of the subject, and changes in both humidity and temperature of the air flowing in and out of the subject with classical cough-like waveforms. Cough detection algorithm is applied with patented fuzzy logic criteria to determine if the event is a cough (64). When analyzing the number of coughs, each cough in a multi-cough event was counted individually.

### Histological assessment of the lung and nasal-associated lymphoid tissue

Upon euthanasia, the rat skull was excised, and the mandible removed. The skulls were fixed in 10% formalin (Fisher Scientific Cat. SF98-4) for 48 hrs at 26 °C. After fixation, formalin was removed, and skulls were frozen at −80 °C until decalcification. Skulls were de-calcified with Richard-Allan Scientific decalcifying solution (Thermo Scientific Ref. 8340-1) at room temperature for 24-48 hrs and embedded in paraffin. Samples were sectioned and stained with Hematoxylin-Eosin (H&E). A Biotek Lionheart Fx was used to scan and image the NALT. ImageJ was used to trace and measure the area of both the left and right NALT using the images from the Lionheart Fx. The left lobe of the lung was excised and fixed in 10% formalin 48 hrs at 26 °C. The left lobe was then embedded in paraffin and stained with H&E by the WVU Pathology Department. H&E stained sections were used to characterize and score acute and chronic inflammation of the lung. All scorings were done by a board-certified pathologist (iHisto). Individual scores were based on a standard qualitative scoring criterion: (0 – none, 1 – minimal (rare), 2 – mild (slight), 3 – moderate, 4 – marked, 5 – severe). Chronic inflammation was characterized by mononuclear infiltrates of the parenchyma, blood vessels, and airway. Acute inflammation scores were assigned due to the presence of neutrophils in the parenchyma, blood vessels, and airway. All examination and scoring were performed blindly as no knowledge of treatment groups were known.

### Imaging of *Bp* in the lung and nasal cavity

Detection of *Bp* in the nasal cavity and the lung were quantified via immunofluorescence (IF) and confocal imaging. The left lobe of the lung and nasal cavity were preserved and sectioned as described above. Sectioned samples underwent deparaffination and rehydration using xylene and ethanol (100%-70%). Antigen retrieval was performed by incubating samples in citrate buffer at 98 °C for 20 min. Samples were blocked using 5% Bovine Serum Albumin (BSA) (Fisher Scientific Cat. 159008) for 1 hr and primarily labeled utilizing a polyclonal rabbit FHA antibody (Gift from Dr. Erik Hewlett) diluted in 1x PBS. Secondary labeling occurred utilizing an anti-rabbit IgG conjugated with Texas Red (Fisher Scientific Cat. T2767) diluted in 1x PBS. Samples were then covered in mounting media (Prolong Gold Antifade reagent with DAPI Cat. 8961). Samples were imaged using a Nikon A1R confocal microscope. Images were analyzed on DAPI channel and at wavelength 650nm for Texas-Red acquisition. Images were acquired using 100x oil immersion lens (100x/1.40 Nikon Plan APO). To identify any potential differences in IF between the two strains, all IF images were deidentified and the microcolonies were manually counted blindly by four volunteers with 2-3 fields of view used per sample.

### Flow cytometry analysis of phagocytes

Neutrophil recruitment in the blood was evaluated by flow cytometry. Blood samples upon collection were then lysed with Pharmylse buffer (BD Biosciences Cat. 555899) for 20 min at room temperature, with slight vortexing throughout. The remaining cells were centrifuged at 1,000 x *g* for 5 min and re-suspended in PBS + 1% fetal bovine serum (FBS) (Gemini Bio Products Cat. 100-500) for 15 min on ice. Samples were then block with anti-CD32 (BD Pharmingen Cat. 550270) antibody for 30 min at 4°C. After incubation, the cells were stained with the appropriate antibody markers; CD45 Alexa flour 700 (Biolegend Cat. 202218), CD161 APC (Biolegend Cat. 205606), CD45R PE Cy7 (eBioscience Cat. 25-0460-82), His48 FITC (eBioscience Cat. 11-0570-82), CD43 PE (Biolegend Cat. 202812), and CD3 VioGreen (Miltenyi Biotec Cat. 130-119-125) (65). After the addition of antibodies, the cells incubated 1 hr at 4°C in the dark. To prepare the lung samples for flow cytometry, the lung homogenate was pushed through a 70 µm cell strainer (BioDesign Cell MicroSives Cat. N70R), creating a single-cell suspension. The suspension was centrifuged at 1,000 x *g* for 5 min. After removal of the supernatant, the pellet was resuspended in Pharmlyse buffer and the cells incubated at 37°C for 2 min. After incubation, the cells were centrifuged at 1,000 x *g* for 5 min, supernatant was removed and blocked and labeled with antibody as described above. The lung and blood samples were centrifuged at 1,000 x *g* for 5 min and the pellets were resuspended in 0.4% paraformaldehyde and stored overnight at 4°C. Samples were washed with 1x PBS and resuspended in 1x PBS for analysis. Cell samples were analyzed on a LSR Fortessa and samples were gated and analyzed using FlowJo v10.

### Lung and serum cytokine and chemokine analysis

Lung homogenate samples were centrifuged at 19,000× *g* for 4 min and the supernatant was removed and stored at −80 °C. Quantitative analysis of cytokines in the serum and lung homogenate was performed using ProcartaPlex Multiplex Immunoassay kit: Cytokine & Chemokine 22-Plex Rat ProcartaPlex^TM^ Panel (Cat. EPX220-30122-901) per the manufacturer’s instructions.

### Serological analysis

Antibody titers of infected rats were measured by ELISA. *Bordetella pertussis* specific ELISA plates were coated with 50 µl of 10^8^ *B. pertussis* grown as mentioned above for infection. For measurement of antibody titers to FHA (Enzo ALX-630-123-0100), PRN (GSK), PT (List Biological Laboratories #180), and ACT (Gift from Dr. Erik Hewlett), we coated plates with 50 µl of each antigen at 1µg/ml. Once coated, plates incubated over night at 4°C. Plates were washed with 1x PBS-Tween 20 and blocked with 5% skim milk for 2 hours at 37°C. Plates were washed again and the serum from the challenge studies were serially diluted down the ELISA plate and incubated for 2 hours at 37°C. After incubation, the plates were washed and coated with 100µl of secondary goat anti-rat IgG (SouternBiotech Cat. 3030-04) and IgM (SouhternBiotech Cat. 3020-04) antibody at a dilution of 1:2,000 in PBS + 5% milk. Once coated, the plate was incubated for 1 hour at 37°C. Plates were then washed again with PBS-Tween 20 and 100 µl *p*- nitrophenyl phosphate substate (Thermo Scientific Cat. 37620) was added, prepared by manufactures instructions, and the plate was developed for 30 min at room temperature for titers to IgG and IgM. Biotek Synergy H1 microplate reader was used to measure the colorimetric signal of the ELISA plate at *A*_450_. Positive antibody titers were determined by any values that were higher than the baseline. The baseline is set as double the average value of the blank, no serum added to these wells. Generating Nightingale rose plots in Python: Data from individual mice were averaged in Microsoft Excel and log transformed. Values were formatted in Excel to be compatible with Python. Data were imported from .csv files using the pandas package and plotted using the “Barpolar” representation feature in the plotly.graph_objects module. Samples with a titer less than 50 were assigned a value of 0.

### Statistical analysis

All data was analyzed using GraphPad Prism 7. The minimum biological replicates for the challenge studies were four. For statistical comparisons between multiple groups over the entire course of the infection a two-way analysis of variance (ANOVA) was used One-way ANOVA was used for comparison between groups for each individual day with Tukey’s post hoc test. Unpaired Student *t*-tests were used for area under the curve (AUC) analysis. Follow-up statistical tests are annotated in figure legends.

### Data availability

Data requests for figures provided can be addressed to the corresponding author.

### Ethics statement

All studies were performed in accordance West Virginia University Institutional Animal Care and Use Committee approved protocol 1811019148.6.

## ACKNOWLEDGMENTS

The preliminary work on this project was in part supported by the Vaccine Development Center at WVU-HSC through a Research Challenge Grant no. HEPC.dsr.18.6 from the Division of Science and Research, WV Higher Education Policy Commission. The project was also supported by NIH R01AI137155 (F.H.D) and CDC Contract (BAA) 75D301-19-R-67835 (F.H.D). Flow cytometry experiments were performed in the West Virginia University Flow Cytometry Core Facility, which is supported by the National Institutes of Health equipment grant number S10OD016165 and the Institutional Development Award (IDeA) from the National Institute of General Medical Sciences of the National Institutes of Health under grant numbers P30GM103488 (CoBRE) and P20GM103434 (INBRE).

JMH, DTB, and JK performed bacterial challenge. JMH, SMK, and JK monitored rat coughing and respiratory function by whole body plethysmography. All authors participated in the animal experiments. JMH and ESK contributed to flow cytometry panel design and analysis. JMH and TYW prepared and ran flow cytometry samples. MAW and MAD performed cytokine analysis. JMH analyzed H&E histology slides. JMH and GJB performed and analyzed immunohistochemistry. JMH performed ELISA assays. JMH, MB, and FHD contributed to experimental design. JMH wrote manuscript with critical revisions from all authors.

The authors would like to thank Dr. Kathleen Brundage (WVU Flow Cytometry & Single Cell Core Facility) for assisting in flow cytometry and equipment instruction. The authors would also like to thank Dr. Amanda Ammer (Microscope Imaging Facility) for support in imaging the IF slides. The authors would also like to thank Dr. James Coad, Jacqueline Karakiozis, and Brice Hickey (Pathology/Histology Core Facility) for the preparation of lung and nasal cavity for IF and performing H&E on the lung and nasal cavity.

The authors would also like to acknowledge that figure 1 was Created with BioRender.com.

